# Multi-omics profiling reveals convergent adaptation of urinary *Pseudomonas aeruginosa* isolates

**DOI:** 10.64898/2026.07.24.740547

**Authors:** Caroline Martin-Duval, Sandrine Dahyot, Abdellah Tebani, Blandine Harel, Erwan Bonnemains, Enzo Debayle, Soumeya Bekri, Jean-Christophe Giard, Martine Pestel-Caron

## Abstract

*Pseudomonas aeruginosa* is a major opportunistic pathogen, responsible for healthcare-associated urinary tract infections. Its metabolic flexibility and genomic plasticity promote its survival and adaptation in complex environments. Here, we conducted an in-depth analysis of three pairs of sequential *P. aeruginosa* urinary isolates, named “early” and “late”, from three patients to investigate metabolic and phenotypic changes during urinary tract adaptation. An integrated multi-omics approach combining RNA sequencing and metabolomics was performed on isolates grown in human urine (HU) and trypticase soy (TS) medium, and compared with previously published proteomics data. Late isolates showed down-regulation of genes encoding type VI secretion system in both media, while oxidative phosphorylation associated-genes were up-regulated in HU. These late isolates also displayed significant down-regulation of amino-acid metabolism suggesting an increased use of carbon sources available in urine. As previously observed in proteomics, siderophore- and iron-related genes were significantly down-regulated for all late isolates in HU but not in TS, supporting convergent adaptation to the low-iron urinary environment. Metabolomic profiles of HU supernatants from late isolates clustered together, showing common metabolite production in HU. The differential metabolic profile between early and late isolates included phosphatidylcholines, acylcarnitines, biogenic amines and amino-acids, highlighting their importance in urinary adaptation. While long-term survival in HU and TS was similar between isolates, biofilm formation was reduced or lost in late isolates in line with the down-regulation of biofilm-related genes. These findings highlight the transcriptomic and metabolic reprogramming as well as the phenotypic changes occurring during *P. aeruginosa* adaptation to the urinary tract.

**Importance:** *Pseudomonas aeruginosa* is a major opportunistic bacterial pathogen responsible for healthcare difficult-to-treat urinary tract infections. Understanding how this bacterium adapts and persists in the human urinary tract is essential for improving the management of persistent infections. Through an integrated analysis of six sequential clinical isolates from three patients, this study shows that *P. aeruginosa* undergoes coordinated metabolic and phenotypic changes that promote survival in the urinary environment. During adaptation, the bacterium reduces traits involved in bacterial competition and biofilm formation while reprogramming its metabolism to the nutrients available in urine. These changes emerged independently in different patients, revealing convergent adaptation to this challenging environment. By identifying biological processes consistently associated with urinary tract persistence, this work provides new insight into the mechanisms that support long-term colonization and highlights adaptive metabolic pathways that may represent future targets for therapeutic intervention.

## Introduction

*Pseudomonas aeruginosa* is a leading cause of nosocomial infections, notably urinary tract infections (UTIs) associated with catheterization, urinary tract abnormalities, and immunocompromised states (1)*. P. aeruginosa* accounts for approximately 10% of catheter-associated UTIs (CAUTIs) (2) and up to 15% of UTIs in intensive care units (3). Due to its ability to form biofilms, particularly on medical devices, and to its high adaptability to various environment, this opportunistic pathogen leads to significant clinical challenges such as frequent relapses and chronic infections (1, 4).

To better understand the mechanisms involved in *P. aeruginosa*’s adaptation to environments encountered in the host, genomic data are important but insufficient to reveal downstream metabolic regulation. Several studies combining multiple omics approaches have analyzed the behavior of *P. aeruginosa* in the lungs of patients with cystic fibrosis (CF), providing specific insights into proteome, transcriptome and metabolome of *P. aeruginosa* in this context (5–7). All these studies described adaptation to the CF environment through metal uptake remodeling, such as a significant increase in iron and other ion transporters, or a decrease in carbon and purine metabolism. They showed that, despite the existence of a shared transcriptional and translational expression pattern, each strain retains a certain degree of flexibility such as significant divergence in metabolomic profiles, particularly in terms of amino-acid variations (5–7).

By contrast, very few global omics studies have investigated *P. aeruginosa* adaptation to the urinary tract and none has integrated proteomic, transcriptomic and metabolomic approaches simultaneously in this context. In a previous work using three pairs of clinical urinary isolates sequentially collected from three patients, we showed that the late isolates with large genomic deletions exhibited slower growth in trypticase soy (TS) and human urine (HU), as well as reduced motility and virulence in the *Galleria mellonella* infection model. Furthermore, proteomic analyses revealed major metabolic rewiring, with opportunism/virulence-related proteins being underabundant in HU-grown late isolates, while energy metabolism-related proteins, such as those involved in the glyoxylate shunt or pentose phosphate pathway, were overabundant in these isolates (8). In particular, siderophore-related proteins were significantly less abundant in late isolates grown in HU, but not in TS, compared to their early counterparts (8). These first results demonstrated that *P. aeruginosa* adapts to the urinary environment according to a specific evolutionary trajectory (8, 9).

In the present study, to gain a comprehensive and dynamic understanding of the intra-patient adaptation of *P. aeruginosa* in urine, we performed global transcriptomic and targeted metabolomic analyses on the three pairs (early and late) of urinary isolates previously characterized by proteomics (8). In addition, long-term survival and ability to form biofilm in HU were assessed.

## Methods

### Bacterial isolates and growth conditions

All experiments were conducted using a panel of six *P. aeruginosa* urinary clinical isolates obtained from three patients (A, D, F) at the Rouen University Hospital (9). In accordance with French law (Article L. 1121-1 of the French Public Health Code), no ethics approval was required, and informed consent was waived for this retrospective, observational study. The study used anonymized microbiological data collected as part of routine clinical practice. This exemption was confirmed by the Ethics Committee for Research on Existing Data and/or Outside the Jardé Law at the Rouen University Hospital, France. Patients characteristics were previously described (9): patients A and D were immunocompromised while patients D and F had urinary tract comorbidities. All three patients had received at least one antibiotic treatment within the six months prior to the first urine collection, and during the period between the two urine samples for the patient A. Each pair was composed of one early (from a first urine sample) and one late (from a subsequent sample and exhibiting large genomic deletions) isolate per patient (9). The nomenclature of the isolates was composed of a letter (A, D and F) which identified the patient, and a small letter (e or l) which corresponded to the early or late isolate, respectively.

Growth was performed in two media: Trypticase Soy broth (TS) and pooled Human Urine (HU) (BioIVT, West Sussex, United Kingdom), as previously described (8). For transcriptomics and metabolomics, isolates were grown overnight in TS at 37°C with shaking (150 rpm) and the inoculum was standardized to an optical density (OD_600nm_) of 0.01 in 10 mL final volume of HU or TS. Cultures were then incubated at 37°C, with shaking at 150 rpm, until late exponential phase. For transcriptomics, cells were centrifuged for 5 min at 13,751 x *g* and washed twice with sterile water to remove the culture medium. The pellets were then resuspended in 200 µL of homogenization solution (Maxwell® RSC Simply Blood Kit, Promega, Wisconsin, USA) and stored at -80°C. For metabolomics, cells were centrifuged for 5 min at 13,751 x *g*. The pellets and supernatants were then stored separately at -80°C.

### RNA extraction

RNAs were extracted from *P. aeruginosa* cells cultivated as described above (late exponential phase) using the Maxwell® RSC simplyRNA Blood Kit (Promega) according to the manufacturer’s instructions. The extracts were quantified with a NanoDrop One spectrophotometer (ThermoFisher Scientific, Massachusetts, USA), and the integrity was assessed using the Agilent TapeStation system (Agilent Technologies, California, USA). Three independent samples for each isolate (A-e, D-e, F-e and A-l, D-l, F-l) and each condition (TS or HU) were prepared.

### RNA-Seq and reads mapping

Independent triplicates of cDNAs from the extracted RNAs (three early and three late isolates grown in HU and TS) were sequenced with a NextSeq 2000 system (Illumina Inc., California, USA) generating 3M reads of 2 x 50 pb (InnovaSEQ platform, University of Caen Normandy, France). cDNA sequences were mapped to the reference genome of *P. aeruginosa* PAO1 (assembly: ASM676v1, accession number: NC_002516.2) using STAR mapper (version 2.7.10b) algorithm (10). Then, a gene count table was created with HTseq-count (version 2.0.9). Normalization and differential expression of RNA counts (late isolate grown in TS/HU for each patient *versus* early isolate grown in TS/HU) were performed using the R© software (v 4.5.2, R foundation, Vienna, Austria, https://www.R-project.org/) and the R package edgeR (version 4) (11). The *p*-value was corrected using the Benjamini–Hochberg method. Gene transcripts with Log2 fold changes (Log2FC) ≥ 2 or ≤ -2 with a corrected *p* value ≤ 0.05 were considered to be significantly deregulated. Metabolic categorization was determined using Kyoto Encyclopedia of Genes and Genomes (KEGG, https://genome.jp/kegg/) and the *Pseudomonas* Genome database (https://www.pseudomonas.com).

### Targeted metabolomic analysis

Three independent samples for each isolate, each medium and each condition (supernatant and pellet, *i.e.* a total of 72 samples) were analyzed with the kit for AbsoluteIDQ® p180 analysis (Biocrates Life Science AG, Innsbruck, Austria). This kit enables the detection of 192 metabolites including 23 amino acids, 23 biogenic amines, 90 phosphatidylcholines, 40 acylcarnitines and 15 sphingomyelins. Sample preparation was performed according to the manufacturer’s protocol as previously described (12). Briefly, 10 µL of sample were loaded in a 96-well plate and dried under a stream of nitrogen. 50 µL of a 5% phenylisothiocyanate (PITC) solution was added to derive biogenic amino acids and amines. After incubation, the spots were dried again before the metabolites were extracted with 5 mM ammonium acetate in methanol (300 µL) in the 96-well plate for analysis after further dilution with the MS running solvent A. Amino acids and biogenic amines were detected in LC-MS mode, while acylcarnitines, phospholipids, sphingomyelins, and the sum of hexoses were analyzed in flow injection analysis (FIA). Quantification was performed using isotopically-labeled internal standards and calibration curve. Analysis was performed using a SCIEX Api4000 QTrap mass spectrometer (AB SCIEX, Darmstadt, Germany) with electrospray ionization. Data acquisition and processing were carried out using the Analyst 1.5 software (Sciex, Framingham, USA). Statistical analyses were performed using R© software. Results were analyzed by an independent *t* test, and comparisons of concentrations with a *p* value of less than 0.05 were considered statistically significant.

### Biofilm formation

Biofilm formation was measured by a crystal violet staining assay. Inoculum of each isolate was prepared from overnight cultures in HU, and standardized to an OD_600nm_ of 0.05. Then, 20 µL of each culture were inoculated into 180 µL of HU in sterile polypropylene microplates (Eppendorf, Montesson, France) in triplicate, and incubated at 37°C for 24h in the Tecan Infinite 200 Pro microplate reader (Tecan, Männedorf, Switzerland), without shaking. Then, OD_595nm_ was measured, and microplates were washes three times with sterile water. Adherent cells were stained with 0.1% (m/v) crystal violet for 10 min, washed and air-dried during 1 h. Biofilms were resuspended with 70% (v/v) ethanol and OD_595nm_ was measured. The *P. aeruginosa* strain PA14 was used as a positive control and fresh medium as a negative control for each experiment. The results represent the mean of absorbance values of three independent experiments. *p* values were determined using independent *t* tests on R© software.

### Long-term survival assays

Each of the six isolates was used to inoculate five independent tubes containing 10 mL of TS or HU. On day 1, each tube was inoculated with 100 µL of an overnight culture in TS at 37°C with shaking at 150 rpm. All tubes were prepared in parallel at the beginning of the experiment but analyzed sequentially (one new tube per week over a 30-day period) to limit the risk of contamination. Tubes were incubated at 37°C, with shaking at 150 rpm. Bacterial survival was assessed every two days by taking a 10 µL aliquot from the corresponding tube. Samples were serially diluted and plated in two technical replicates on TS agar plates. After 24 hours of incubation at 37°C, CFUs were counted. All experiments were repeated at least three times. Statistical analyses were performed using independent *t* tests. Differences with a *p* value ≤ 0.05 were considered statistically significant.

## Results

### Transcriptional convergence of *P. aeruginosa* in urinary isolates

#### Overall comparison of gene expression between early and late isolates

The global transcriptomic study of the three pairs of early and late isolates from patients A, D and F grown in TS and HU media was performed to identify putative transcriptional signature associated with *P. aeruginosa* adaptation to the urinary tract. A total of 931 (661 upregulated and 270 downregulated) and 2,039 (855 upregulated and 1,184 downregulated) genes were differentially expressed when comparing all three late isolates grown in TS and in HU respectively, with their early counterparts (**Fig. 1, Table I**). For patients D and F, HU-grown late isolates displayed significantly more down-regulated genes (*n* = 435 and 462, respectively) than up-regulated genes (*n* = 176 and 188, respectively) (**Table I**). The opposite was observed for the A-l isolate grown in HU (491 up-regulated and 287 down-regulated genes) (**Table I**). It should be noted that the twelve most up-regulated genes for D-l and F-l isolates grown in TS and HU (from +6.42 to +12.3 logFC), belonged to the same *Alg* operon (**Table S2**), responsible for alginate biosynthesis, a major exopolysaccharide in this species (13). Conversely, the most strongly down-regulated genes in late isolates belonged to two of the three operons encoding the type VI secretion system (T6SS) (up to -7.13 logFC in HU and up to -5.11 logFC in TS), namely H1-T6SS and H3-T6SS, with some of the H2-T6SS genes, also being down-regulated (**Table S2**). For example, for the A-l isolate grown in HU, 18 out of 34 genes belonging to the H1-T6SS operon were down-regulated.

**FIG 1.**
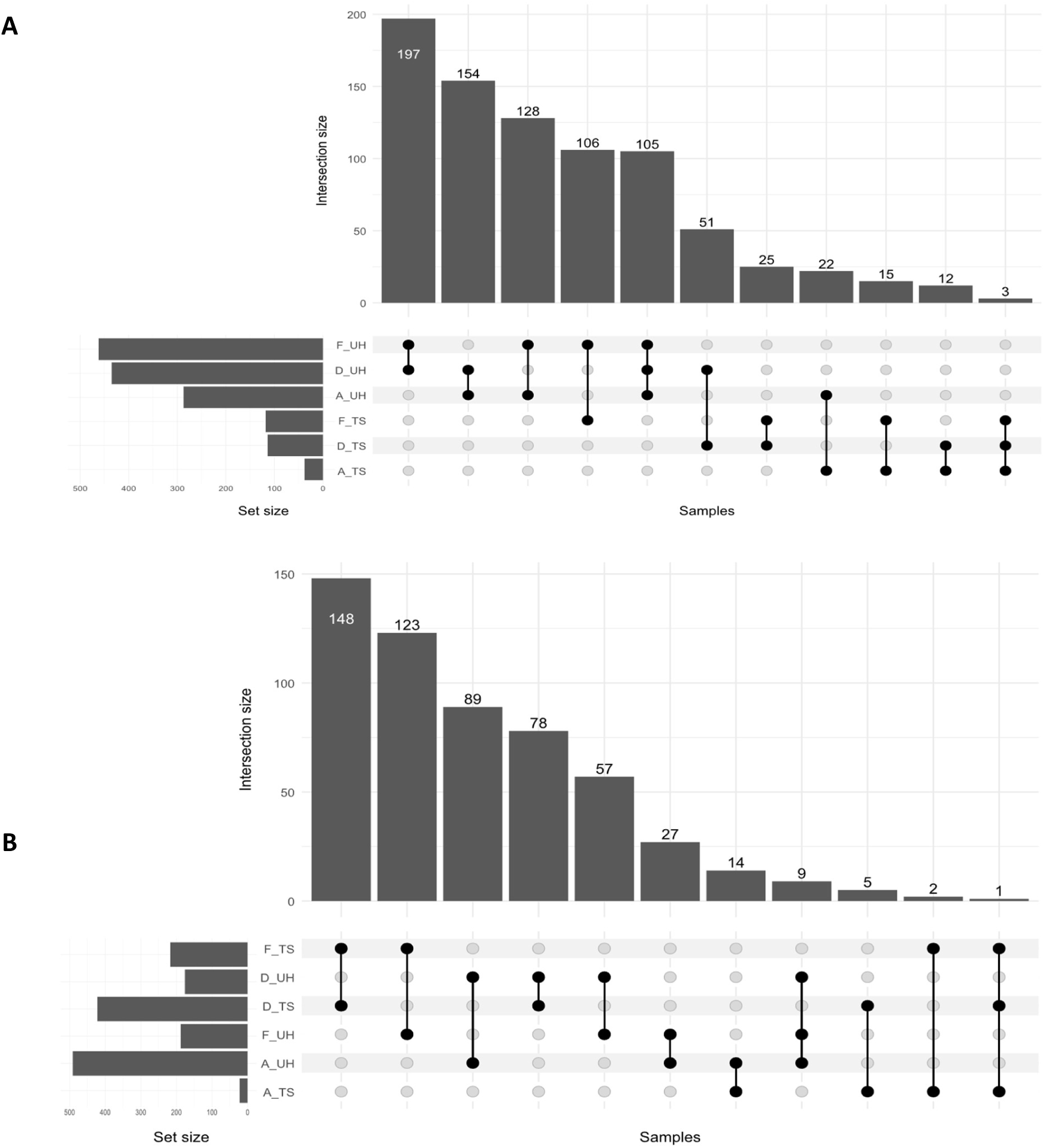
Down-regulated (A) and up-regulated (B) genes across early and late isolates from patients A, D and F in two conditions (Trypticase Soy broth [TS] or pooled Human Urine [HU]). The bar chart at the top shows the intersection size of each combination. The bar chart on the left displays the number of genes in each condition. The dot matrix at the bottom represents individual gene sets for each patient-condition combination, and connected black dots represent shared gene sets across multiple conditions.

**Table I.**
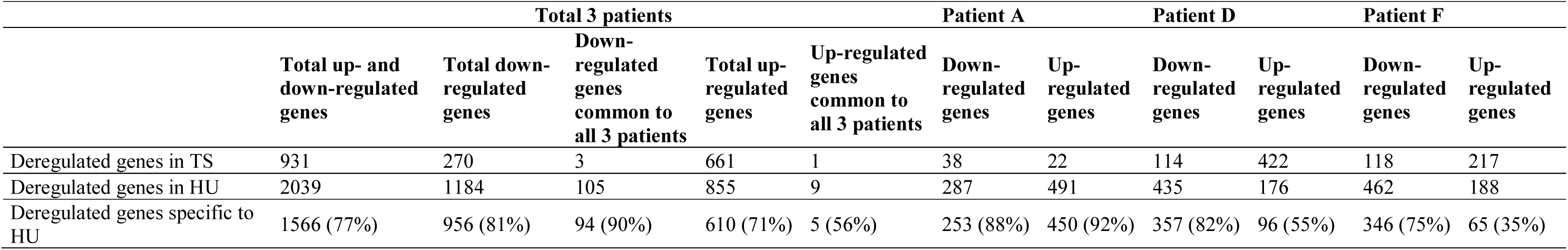
Number of down-regulated or up-regulated genes in late isolates compared to early isolates in Trypticase Soy (TS) and Human Urine (HU). The number of HU-specific deregulated genes is specified.

|  | Total 3 patients |  |  |  |  | Patient A |  | Patient D |  | Patient F |  |
| --- | --- | --- | --- | --- | --- | --- | --- | --- | --- | --- | --- |
|  | Total up- and down-regulated genes | Total down-regulated genes | Down-regulated genes common to all 3 patients | Total up-regulated genes | Up-regulated genes common to all 3 patients | Down-regulated genes | Up-regulated genes | Down-regulated genes | Up-regulated genes | Down-regulated genes | Up-regulated genes |
| Deregulated genes in TS | 931 | 270 | 3 | 661 | 1 | 38 | 22 | 114 | 422 | 118 | 217 |
| Deregulated genes in HU | 2039 | 1184 | 105 | 855 | 9 | 287 | 491 | 435 | 176 | 462 | 188 |
| Deregulated genes specific to HU | 1566 (77%) | 956 (81%) | 94 (90%) | 610 (71%) | 5 (56%) | 253 (88%) | 450 (92%) | 357 (82%) | 96 (55%) | 346 (75%) | 65 (35%) |

#### Genes and corresponding pathways up-regulated in late isolates

As shown in the **Fig. 2**, the functional categories with the highest number of up-regulated genes in HU-grown late isolates were linked to cellular adaptation and energy metabolism. Indeed, 36% (*n* = 178/491) and 26% (*n* = 45/176) of up-regulated genes were involved in translation in late isolates of A and D respectively, represented for a large part by ribosome-related genes. Of note, this gene category was underrepresented in F-l isolate with only one amino-acyl-tRNA biosynthesis gene among the 188 upregulated genes. The cellular adaptation genes over-transcribed in F-l were instead involved in alginate biosynthesis (with six genes linked to fructose and mannose metabolism) and six genes involved in exopolysaccharide biosynthesis. Regarding energy metabolism, oxidative phosphorylation was the category in which a large number of genes were up-regulated for all late isolates from patients A, D and F grown in HU (*n* = 30/491, 10/176 and 14/188 respectively) (**Fig. 2, Table S2**).

**FIG 2.**
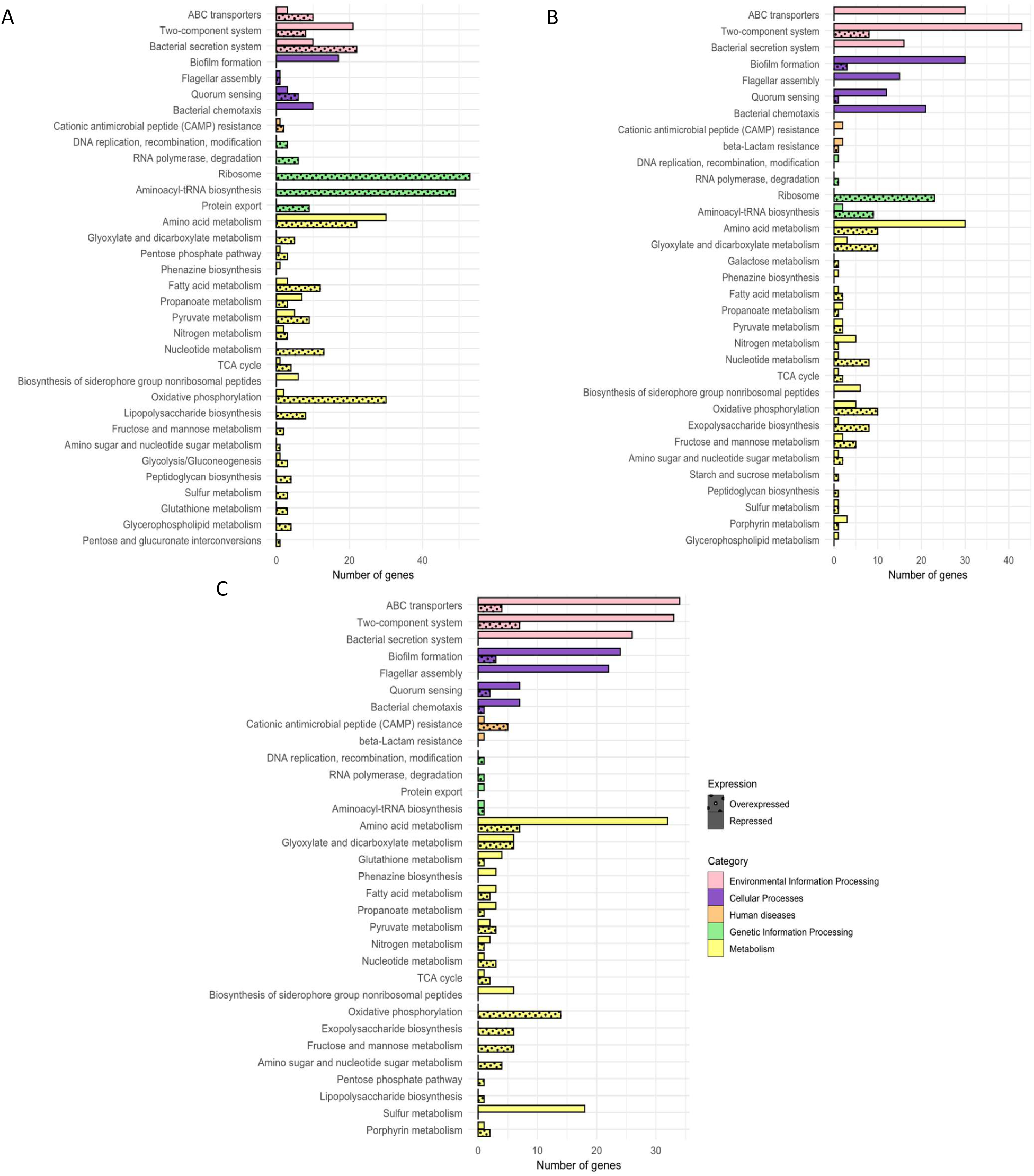
Functional classification according to KEGG pathway of the significantly down-regulated (full bars) and up-regulated (dotted bars) genes of late isolates from patients A (A), D (B) and F (C) grown in human urine compared to early isolates. Only genes with a log2 fold change (log2FC) ≥ 2 and ≤ -2 were considered. Pathways are color-coded by major functional categories: environmental information processing (pink), cellular processes (purple), human diseases (orange), genetic information processing (green) and metabolism (yellow).

Only nine up-regulated genes were common to the three late isolates, whatever the medium (TS and HU) (**Table S1**). Seven of them were involved in energy metabolism, coding GDP-mannose 6-dehydrogenase AlgD, a 2,3 butanediol dehydrogenase, two nitrate-inducible formate dehydrogenases FdnH and FdnI (beta and gamma subunit respectively), an ATP-dependent RNA helicase DeaD and a cbb3-type cytochrome c oxidase subunit CcoN1. The proteins encoded by these genes play a role in alginate biosynthesis, butanoate metabolism, glyoxylate cycle, RNA metabolism and oxidative phosphorylation respectively (14–18). Five common up-regulated genes appeared specific to the growth in urine and concerned energy metabolism including the glyoxylate and dicarboxylate pathways and the oxidative phosphorylation metabolism.

#### Genes and corresponding pathways down-regulated in late isolates

Genes that were downregulated in late isolates grown in HU belonged mainly to the amino acid metabolism pathway (*n* = 30/287, 30/435 and 34/462 genes for A-l, D-l and F-l, respectively), two-component systems as *fleS/R* involved in flagellum-dependent motility or *foxI/R* involved in iron uptake, but were also related to biofilm formation (*i.e. psl* and *rhl* genes) (**Fig. 2, Table S2**). Moreover, for patients D and F, a substantial number of ABC transporters-related genes were also downregulated (*n* = 30/435 and 34/462 for D-l and F-l, respectively).

Ninety-four down-regulated genes were observed in all late isolates grown in HU but not in TS. These were mostly iron-related genes involved in pyoverdine and pyochelin biosynthesis (the two main siderophores of *P. aeruginosa*) or heme-related genes (60%, *n* = 56/94), that are important to survive in iron-limited environments. 105 of the down-regulated genes observed in all HU-grown late isolates were also found when these cells were grown in TS (**Table S1**). They were mainly involved in virulence such as biofilm formation, iron regulation/transport or the type VI bacterial secretion system as well as in amino acid biosynthesis. Of note, genes encoding subunits of the T6SS corresponded to a large proportion of the down-regulated genes (from 6% (*n* = 29/462) for F-l in HU to 53% (*n* = 16/30 for A-l in TS) (**Fig. 2, Table S2**).

## Transcriptome correlates with proteome data

We then compared transcriptomic results with locus tag previously identified by proteomics analyzed under the same *in vitro* conditions (8). A correspondence of up to 62% (87 up-regulated genes out of 141 overabundant proteins for D-l) was observed in TS condition, and up to 43% (45 up-regulated genes out of 105 overabundant proteins for D-l) in HU (**Table S2,** EMBL-EBI accession number PRJEB114131). Moreover, 27 locus tags downregulated were found in common in both transcriptomics and proteomics for the three late isolates grown in HU (**Table II**) while no upregulated locus tags common to both approaches were identified.

**Table II.**
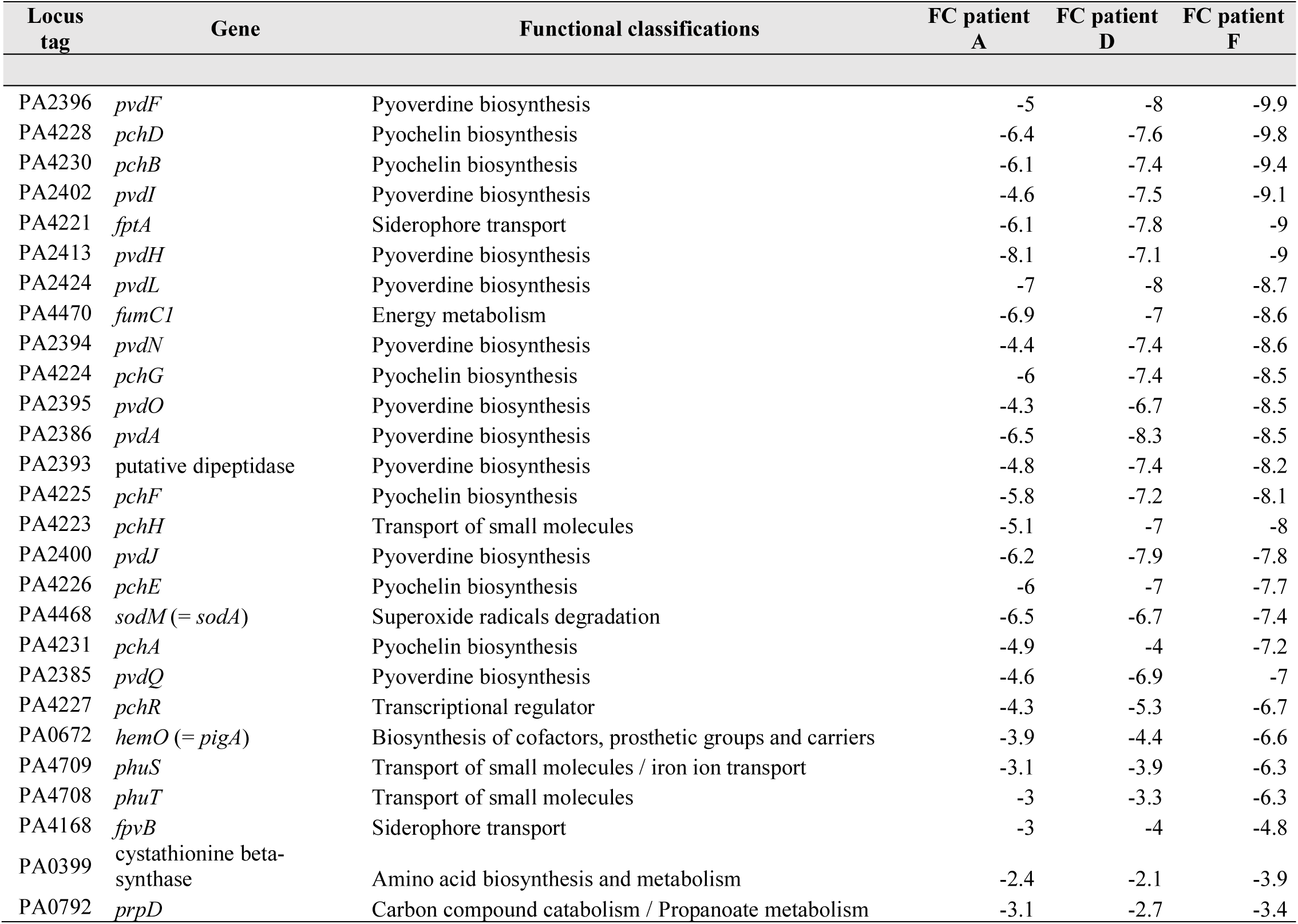
List of the 27 locus tags significantly down-regulated in both transcriptomics and proteomics (8) in the three late isolates (from patients A, D and F) grown in Human Urine (HU). Gene functional classification was performed using the *Pseudomonas* Genome Database.

| Locus tag | Gene | Functional classifications | FC patient A | FC patient D | FC patient F |
| --- | --- | --- | --- | --- | --- |
| PA2396 | *pvdF* | Pyoverdine biosynthesis | -5 | -8 | -9.9 |
| PA4228 | *pchD* | Pyochelin biosynthesis | -6.4 | -7.6 | -9.8 |
| PA4230 | *pchB* | Pyochelin biosynthesis | -6.1 | -7.4 | -9.4 |
| PA2402 | *pvdI* | Pyoverdine biosynthesis | -4.6 | -7.5 | -9.1 |
| PA4221 | *fptA* | Siderophore transport | -6.1 | -7.8 | -9 |
| PA2413 | *pvdH* | Pyoverdine biosynthesis | -8.1 | -7.1 | -9 |
| PA2424 | *pvdL* | Pyoverdine biosynthesis | -7 | -8 | -8.7 |
| PA4470 | *fumC1* | Energy metabolism | -6.9 | -7 | -8.6 |
| PA2394 | *pvdN* | Pyoverdine biosynthesis | -4.4 | -7.4 | -8.6 |
| PA4224 | *pchG* | Pyochelin biosynthesis | -6 | -7.4 | -8.5 |
| PA2395 | *pvdO* | Pyoverdine biosynthesis | -4.3 | -6.7 | -8.5 |
| PA2386 | *pvdA* | Pyoverdine biosynthesis | -6.5 | -8.3 | -8.5 |
| PA2393 | putative dipeptidase | Pyoverdine biosynthesis | -4.8 | -7.4 | -8.2 |
| PA4225 | *pchF* | Pyochelin biosynthesis | -5.8 | -7.2 | -8.1 |
| PA4223 | *pchH* | Transport of small molecules | -5.1 | -7 | -8 |
| PA2400 | *pvdJ* | Pyoverdine biosynthesis | -6.2 | -7.9 | -7.8 |
| PA4226 | *pchE* | Pyochelin biosynthesis | -6 | -7 | -7.7 |
| PA4468 | *sodM* (= *sodA*) | Superoxide radicals degradation | -6.5 | -6.7 | -7.4 |
| PA4231 | *pchA* | Pyochelin biosynthesis | -4.9 | -4 | -7.2 |
| PA2385 | *pvdQ* | Pyoverdine biosynthesis | -4.6 | -6.9 | -7 |
| PA4227 | *pchR* | Transcriptional regulator | -4.3 | -5.3 | -6.7 |
| PA0672 | *hemO* (= *pigA*) | Biosynthesis of cofactors, prosthetic groups and carriers | -3.9 | -4.4 | -6.6 |
| PA4709 | *phuS* | Transport of small molecules / iron ion transport | -3.1 | -3.9 | -6.3 |
| PA4708 | *phuT* | Transport of small molecules | -3 | -3.3 | -6.3 |
| PA4168 | *fpvB* | Siderophore transport | -3 | -4 | -4.8 |
| PA0399 | cystathionine beta-synthase | Amino acid biosynthesis and metabolism | -2.4 | -2.1 | -3.9 |
| PA0792 | *prpD* | Carbon compound catabolism / Propanoate metabolism | -3.1 | -2.7 | -3.4 |

These 27 down-regulated locus tags belonged to two main metabolic pathways: iron metabolism and carbon metabolism (**Table II**). Indeed, 85% (*n* = 23/27) were iron-related genes or proteins, and more specifically, ten were involved in pyoverdine biosynthesis (43%) and eight in pyochelin biosynthesis (35%). Three genes involved in pyoverdine biosynthesis were organized in a single operon, while seven others associated with pyochelin biosynthesis were distributed across two separate operons (19). In addition, expression of two main ferric-siderophore receptors were also down-regulated : *fptA* specific for pyochelin and *fpvB* for pyoverdine (20). Three loci were involved in carbon metabolism: PA0399, encoding a cystathionine beta-synthase; *fumC1* encoding a fumarate hydratase involved in the tricarboxylic acid cycle and redox balance (21), and *prpD* encoding a propionate catabolic protein, involved in the propanoate metabolism (**Table II**). Note that in the same operon as *fumC1*, the *sodM* gene, encoding a superoxide dismutase involved in the oxidative stress response, was also down-regulated.

## Metabolomic profiles

To better understand metabolic adaptation, we expanded our approach by integrating a targeted metabolomic analysis of early and late isolates grown in TS and HU (cell pellets and supernatants). We analyzed 192 metabolites including lipids (phosphatidylcholines, acylcarnitines, sphingomyelins), biogenic amines and amino acids. An unsupervised analysis was carried out using metabolomic data as previously described (22). Principal component analysis (PCA) score plots are presented in **Fig. 3**.

**FIG 3.**
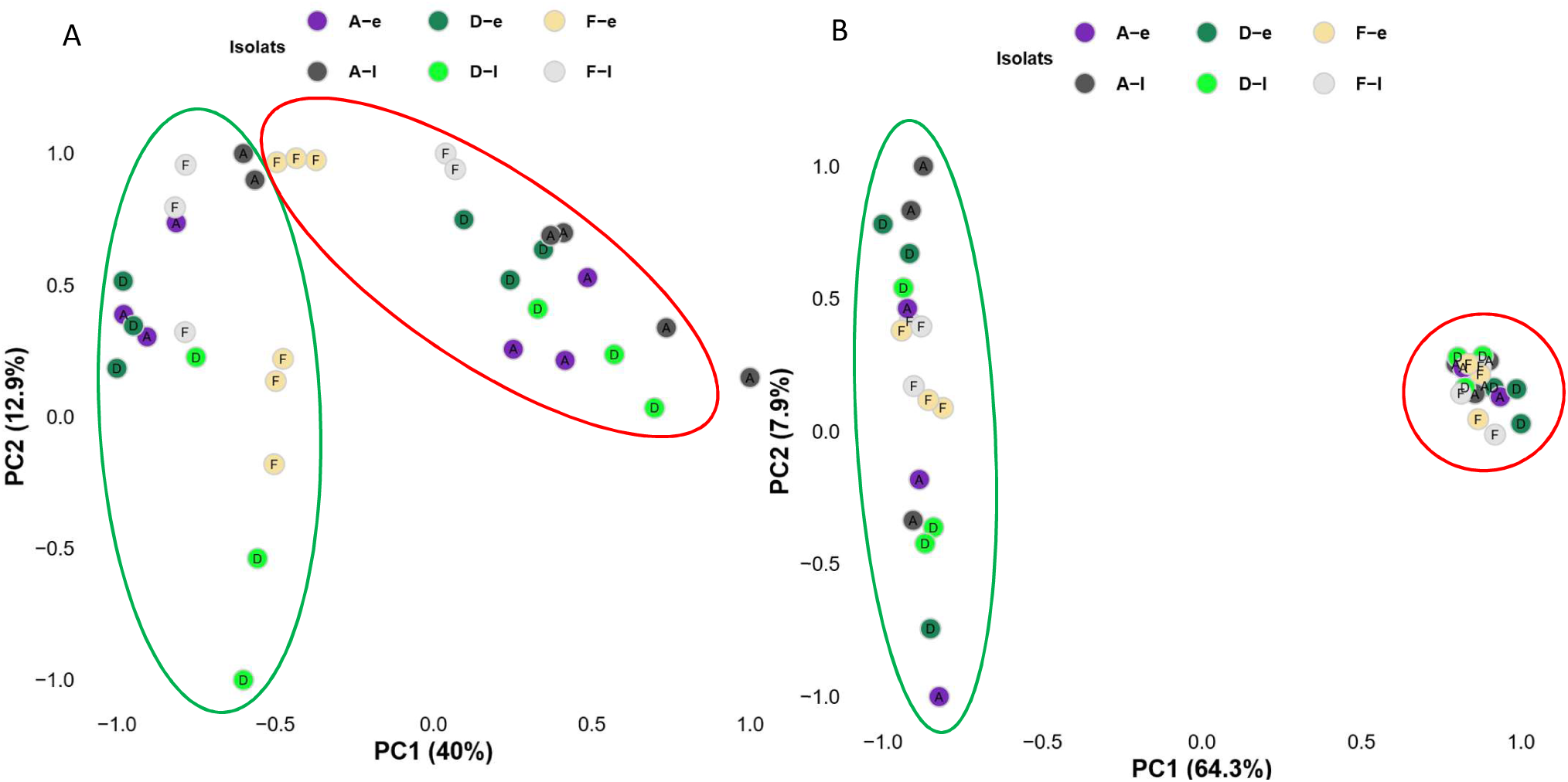
Principal component analysis score plots of the metabolites identified in cell pellets (A) and supernatants (B) of early and late isolates from patients A, D and F grown in TS (circled in green) and HU (circled in red) in three independent experiments.

These PCA showed that the variance in metabolites from bacterial pellets and supernatants grown in HU is mainly explained by the PC1 (40% and 64.3% respectively) (**Fig. 3**). This highlights a metabolic shift echoing adaptation of the isolates in their environment. Moreover, the sample dispersion from the supernatants of the bacterial cells grown in urine was markedly lower than that of samples from the pellets. On the other hand, in TS, the dispersion of samples on the PCA scores was as heterogeneous from the pellet samples as it was from the supernatant samples (**Fig. 3**). This may reflect a greater diversity of the metabolite production by the early and late isolates in the rich medium compared to HU (**Fig. 3**).

In cell pellets, 24 of the 192 metabolites were detected when isolates grown in TS and 23 in HU medium. 35% of amino acids (AAs) (*n* = 8), 13% of biogenic amines (BAs) (*n* = 3) and 13% of phosphatidylcholines (PCs) (*n* = 12) were detected in the cell pellets cultivated in TS and HU media. More metabolites were detected in supernatants, with 41 of 192 in TS cultures and 39 in HU cultures. In the supernatants of HU cultures, 48% were AAs (*n* = 11), 48% ACs (*n* = 19), 30% BAs (*n* = 7) and 2% PCs (*n* = 2), while in TS supernatants, 65% were AAs (*n* = 15), 45% ACs (*n* = 18), 30% BAs (*n* = 7), and 1% PCs (*n* = 1).

Metabolites whose concentrations were significantly different between cultures of early and late isolates in HU or TS (*p* < 0.05) were summarized in **Table III**. No common metabolic profile was observed among the late isolates for PCs and ACs. Of note, for HU-grown isolates, alanine was identified in significantly lower concentrations in the A-l pellet and in significantly higher concentrations in the D-l and F-l supernatants compared to early isolates (**Table III**). In contrast, glutamate was detected in significantly lower concentrations in the D-l and F-l pellets and in significantly higher concentrations in the F-l supernatant compared to early isolates (**Table III**). PC 32:0 was the only metabolite measured at a significantly higher concentration in the D-l cell pellet compared to that of D-e isolate in both HU and TS media (*p* < 0.05) (**Table III**).

**Table III.**
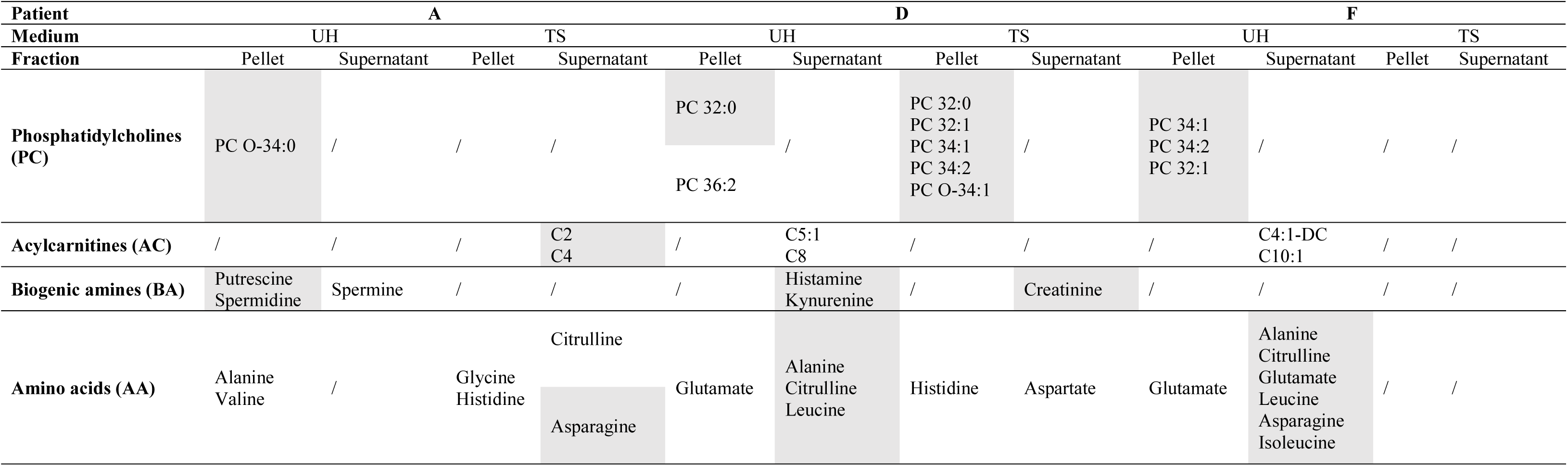
Metabolites showing a significant concentration difference between early and late isolates from patients A, D and F grown in human urine (HU) or in trypticase soy (TS) as part of a targeted metabolomic analysis on cell and pellet samples. The grey boxes represent metabolites whose concentrations were significantly higher for the late isolate than for the early isolate, and conversely for the white boxes. Statistical significance was determined as *p* ≤ 0.05 using a two-sample Student’s *t*-test for independent samples (*n* = 3).

| Patient | A |  |  |  | D |  |  |  | F |  |  |  |
| --- | --- | --- | --- | --- | --- | --- | --- | --- | --- | --- | --- | --- |
| Medium | UH |  | TS |  | UH |  | TS |  | UH |  | TS |  |
| Fraction | Pellet | Supernatant | Pellet | Supernatant | Pellet | Supernatant | Pellet | Supernatant | Pellet | Supernatant | Pellet | Supernatant |
| Phosphatidylcholines (PC) | PC O-34:0 | / | / | / | PC 32:0 | / | PC 32:0<br>PC 32:1<br>PC 34:1<br>PC 34:2<br>PC O-34:1 | / | PC 34:1<br>PC 34:2<br>PC 32:1 | / | / | / |
|  |  |  |  |  | PC 36:2 |  |  |  |  |  |  |  |
| Acylcarnitines (AC) | / | / | / | C2<br>C4 | / | C5:1<br>C8 | / | / | / | C4:1-DC<br>C10:1 | / | / |
| Biogenic amines (BA) | Putrescine<br>Spermidine | Spermine | / | / | / | Histamine<br>Kynurenine | / | Creatinine | / | / | / | / |
| Amino acids (AA) | Alanine<br>Valine | / | Glycine<br>Histidine | Citrulline | Glutamate | Alanine<br>Citrulline<br>Leucine | Histidine | Aspartate | Glutamate | Alanine<br>Citrulline<br>Glutamate<br>Leucine<br>Asparagine<br>Isoleucine | / | / |
|  |  |  |  | Asparagine |  |  |  |  |  |  |  |  |

## Biofilm formation

As transcriptomics showed the down-regulation of numerous genes involved in biofilm formation for late isolates (**Fig. 2**), the ability to form such a structure in urine was explored (**Fig. 4**). The early isolates from the three patients showed a significantly higher biofilm forming capacity than the reference strain PA14 (*p* = 0.0097, *p* = 1.1×10^-6^, *p* = 0.025 respectively). Otherwise, no biofilm formation was observed for A-l and F-l isolates, whereas a 15-fold reduction in production was measured for D-l compared to D-e (*p* = 7.94×10^-7^) (**Fig. 4**).

**FIG 4.**
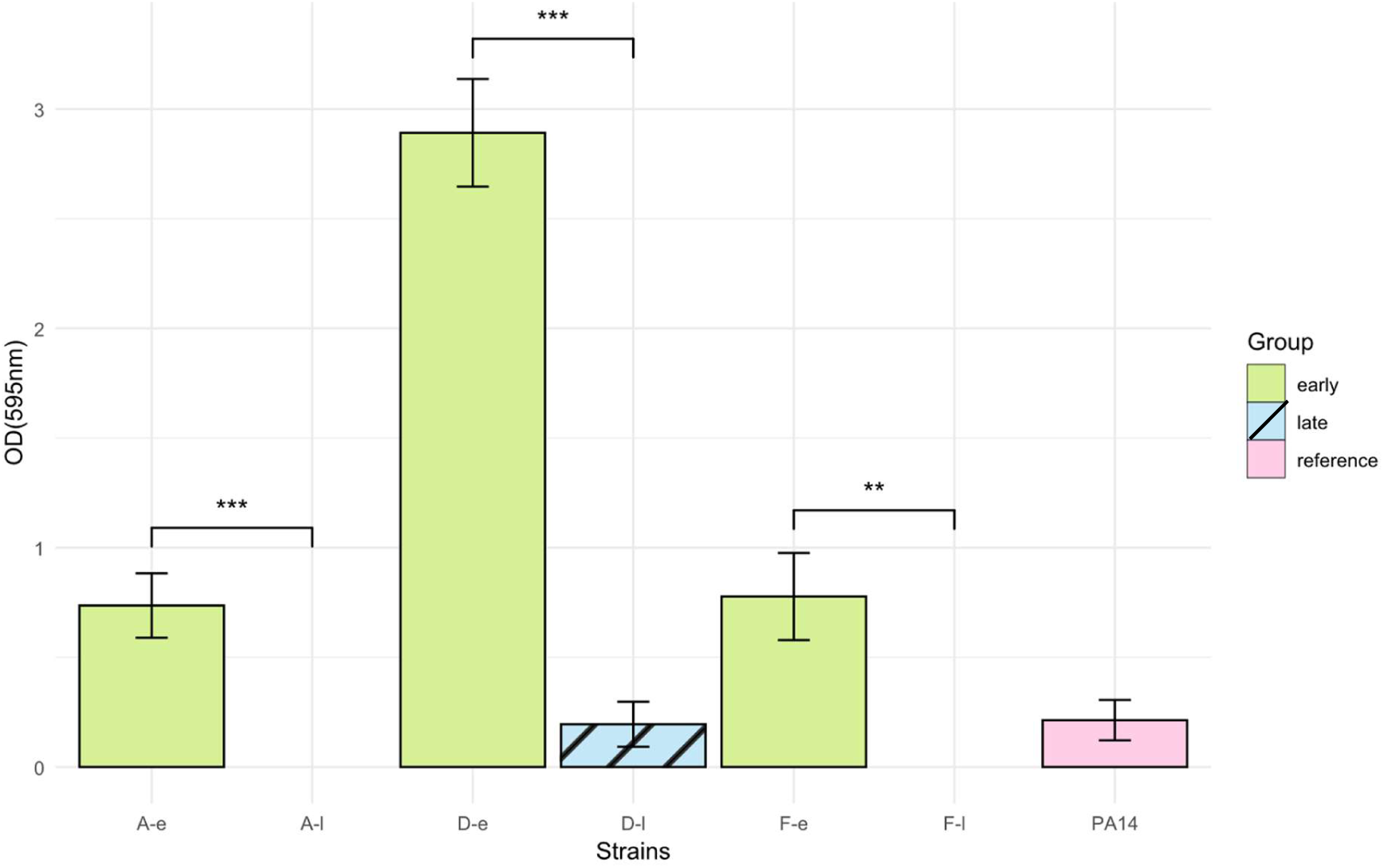
Biofilm formation of early and late isolates from patients A, D and F when cultured in HU. Error bars represent the standard deviations of three independent experiments. Statistical significance was determined as *p* ≤ 0.05 using a two-sample Student’s *t*-test for independent samples.

Overall, these results were consistent with the transcriptomic and proteomic observations as 17, 30 and 24 biofilm-related genes and 10, 8 and 5 biofilm-related proteins were down-regulated and underabundant in A-l, D-l and F-l HU-grown isolates respectively, compared with their early counterparts (8).

## One-month survival assay

To test the hypothesis that late isolates have better survival capacity than early ones, long-term survival experiments were conducted for 30 days in HU and TS media for each isolate (**Fig. 5**).

**FIG 5.**
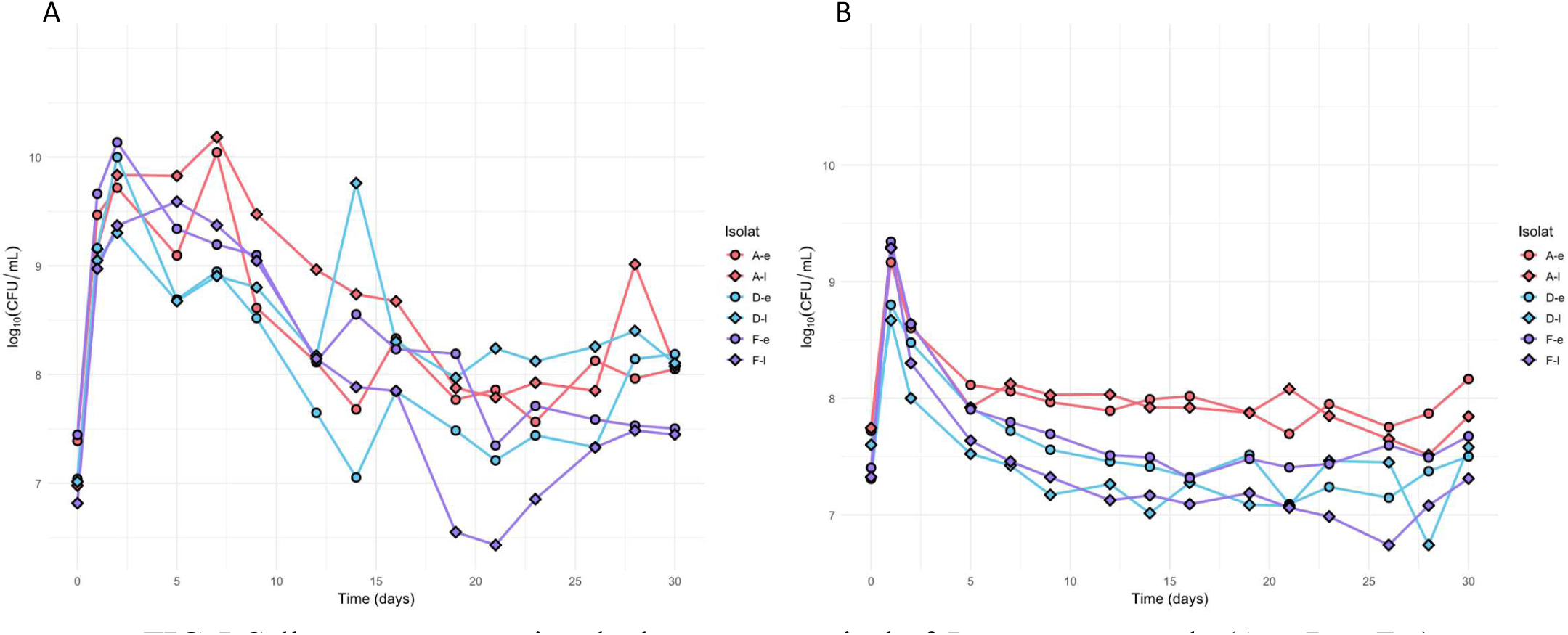
Cell counts representing the long-term survival of *P. aeruginosa* early (A-e, D-e, F-e) and late (A-l, D-l, F-l) isolates in Trypticase Soy broth (A) or in Human Urine (B).

In TS medium, the late isolates from patients A and D displayed a significantly higher CFU/mL counts than early isolates at different time points (*p* < 0.05) (**Fig. 5A**). However, for isolates from patient F, a significantly higher number of CFU/mL for F-e was observed only between day 1 and day 5 (*p* < 0.05) (**Fig. 5A**). Conversely, in HU medium, all early isolates presented a significant higher CFU/mL count than their late counterparts at several time points (**Fig. 5B**). For example, A-e had significantly better survival at day 28 (A-e = 7.4.10^7^ CFU/mL; A-l = 3.3×10^7^) and day 30 (A-e = 1.5×10^8^ CFU/mL, A-l = 7.0×10^7^ CFU/mL) than A-l (*p* < 0.05) (**Fig. 5B**). Taken together, these results revealed a variable survival of early and late isolates in urine, with light but significant higher bacterial count for early isolates at different time points. After 30 days in urine, despite a nutrient depletion, CFU/mL counts remained important (≈ 10^7^ CFU/mL) whatever the isolate.

## Discussion

*P. aeruginosa* is well-known for its genomic and metabolic versatility, which allows it to colonize a wide range of ecological niches. Despite its involvement in healthcare-associated UTIs (2), very few data are available on *P. aeruginosa* survival strategies in urine, especially through multi-omic approaches. However, understanding *P. aeruginosa* adaptive mechanisms in the urinary tract could pave the way for new therapeutic and preventive strategies to treat recurrent UTI caused by this ESKAPE pathogen (23).

This study follows on from previous work on the phenotypic, genomic and proteomic characterization of sequential urinary isolates of *P. aeruginosa* (8, 9). Our objective was to improve our understanding of *P. aeruginosa* urinary pathoadaptation, by supplementing our previous data with an original approach combining transcriptomic and targeted metabolomic analyses.

First, late isolates up-regulated genes involved in oxidative phosphorylation. This suggests adaptation to a microaerophilic environment (16) and a need for optimization of oxidative phosphorylation. In addition, among the most up-regulated genes were those belonging to the *alg* operon, responsible for alginate biosynthesis, a major exopolysaccharide in this species (13). This could partially explain the mucoid morphotype of the F-l isolate (9). By contrast, Tielen and collaborators reported a decrease in alginate formation under urinary-tract like conditions (24). However, it should be noted that this work used the PAO1 reference strain (isolated in 1954 from a wound) grown anaerobically in artificial urine medium (24). This highlights the importance of working with clinical urinary isolates and pooled human urine, to replicate physiological conditions as closely as possible. It is interesting to note that only five genes were commonly up-regulated in the three late isolates grown in HU, mainly involved in energy metabolism, including oxidative phosphorylation and the glyoxylate and dicarboxylate pathway. The latter reflects an adaptation of the isolates’ metabolism to the host and environment through more efficient nutrient assimilation (25).

On the other hand, genes involved in virulence, with siderophore biosynthesis, secretion systems, and flagellar assembly, were down-regulated in late isolates, as previously demonstrated in proteomics (8). These observations showed gene-protein regulation in the same direction and correlated with the less virulent phenotype of late isolates previously identified in the *Galleria mellonella* larval infection model (8). These characteristics have already been identified in CF adapted isolates as a convergent evolutive trajectory (26, 27). Indeed, various studies have shown an accumulation of nonsynonymous mutations in motility and virulence genes (26), and a decrease in lethality in a murine model of chronic airways infection (27). Interestingly, numerous iron-related genes were also repressed, in correlation with the underabundance of the corresponding proteins in HU-grown late isolates (8), despite the fact that urine is an iron-limited medium (28). This indicates a broad and coordinated decrease in iron acquisition pathways in these isolates. This regulatory pattern is not consistent with previous work on cystic fibrosis or CAUTI (5, 29), where iron is also poorly available. This suggests a specific adaptive response of these urinary isolates which reduce their dependence on iron to decrease their energy consumption, as siderophores are very costly to synthesize (30). Among the down-regulated genes related to virulence, up to 30 genes were involved in biofilm formation in HU including the *psl* and *rhl* genes (associated with underabundance of the corresponding proteins), which could explain the drastic reduction in biofilm formation observed for late isolates. These data are in contrast with those described in CF lungs or in CAUTI, where epithelial and catheter colonization by *P. aeruginosa* is associated with the biofilm development (31–33).

Transcriptomic and proteomic (8) approaches also revealed a down-regulation of respectively genes and proteins involved in amino acid metabolism, common to all late isolates grown in HU, particularly those related to phenylalanine, tyrosine and tryptophan. These amino acids are scarce in urine (28) and are less assimilated by *P. aeruginosa* than other amino acids or short-chain fatty acids (34). Adaptation in amino acid metabolism has already been demonstrated for late isolates of *P. aeruginosa* collected from CF lungs (35, 36). By exometabolome analyses, La Rosa and coworkers identified a nutritional hierarchy in terms of amino acid assimilation, particularly depending on the growth phase and limiting amino acid biosynthesis (36).

To further characterize metabolic specialization of *P. aeruginosa* in urine, targeted metabolomic analyses looking for phosphatidylcholines, acylcarnitines, sphingomyelins, biogenic amines and amino acids were performed. These analyses were performed in both cell pellets and supernatants of each isolate to characterize intracellular and extracellular metabolic changes between early and late isolates. Metabolomic data supported transcriptomic and proteomic observations. First, the PCA score plots showed that the supernatants of HU-grown samples formed a tight cluster, unlike those grown in the rich TS medium. It is important to note that this clustering reflects homogeneity in the bacterial metabolomic profiles identified, and not the chemical complexity of the growth media, as human urine is inherently more complex than TS (37). This illustrates the metabolic convergence of *P. aeruginosa* isolates in urine, which use and produce metabolites in a similar way. It would have been interesting to include a non-urinary *P. aeruginosa* isolate (*e.g.* from a cystic fibrosis patient) cultured in HU, in order to compare this isolate with our urinary isolates, thereby determining whether the clustering observed is due to the origin of the isolate or its adaptation to the urinary environment.

Then, metabolite analysis revealed that supernatants from late isolates grown in HU showed an enrichment in amino acids compared to early isolates, consistent with the reduced expression of genes and proteins involved in this metabolism identified by transcriptomics and proteomics. This suggests a decrease in endogenous amino acid biosynthesis, and an increased dependence on amino acids present in urine, albeit at low concentrations (28).

There was no convergent profile among the late isolates, except some similarities between those of D-l and F-l isolates grown in HU which could reflect the convergence also found in transcriptomics and proteomics. This suggests that these two isolates were probably better adapted to the urinary environment than A-l isolate, which would be correlated with the fact that the infection or colonization in patients D and F were older than in patient A (data not available).

Metabolite profiles suggested the use of BAs and lipids (PCs or ACs) as alternative nutrient sources to survive in urine, consistent with previously observations in *P. aeruginosa* during cystic fibrosis lung infection (38). The increase in PCs (compounds that can be metabolized into glycine-betaine, a well-known osmoprotectant), in D-l and F-l pellets could be evidence of the catabolism of these lipids in response to the osmotic stress present in urine (39, 40). This can be correlated with the proteomic overabundance of two ABC transporters OpuCA and OpuCD, involved in the induction of the hyperosmotic salinity response, identified in late isolates (8). Interestingly, no evidence of PCs catabolism was shown despite a pellet enrichment, suggesting an incorporation in cell membranes, has already described in *P. aeruginosa* CF isolates (41).

The survival of late isolates for 30 days, despite conditions that do not accurately reflect the physiological renewal of urine in the bladder, suggests a resource-optimization strategy resulting in a reorganization of metabolism based on available substrates as has already been described for this species (34). However, to strengthen our conclusions, it would be necessary to repeat the multi-omic analyses and phenotypic experiments after 30 days of survival, under more physiological conditions, for example by renewing the medium daily.

By combining transcriptomic and metabolomic approaches with phenotypic characterization, we provided a comprehensive view of *P. aeruginosa* adaptation to urine, which reinforced our previous assumptions (8). Our findings support the idea that chronic urinary isolates adopt fine adaptative persistence strategy. They reorganize metabolism through available nutrients and downregulate energy-expensive or unnecessary traits to reduce metabolic burden, as already described for CF isolates (42). Furthermore, they drastically reduce their pathogenicity, probably to remain undetectable by the immune system (43). Importantly, this work sheds light on possible preventive and therapeutic approaches strategies targeting bacterial metabolism and nutrient acquisition pathways, in line with the concept of nutritional immunity (44). As mentioned by Bontemps-Gallo in its published comment about our previous work, these findings again highlighted the “cost of chronicity as an adaptive trade-off between acute infection and long-term colonization” (45). Future studies should extend these analyses to larger collections of isolates, including as many sequential clinical isolates as possible, in order to better illustrate the dynamics of urinary pathoadaptation of *P. aeruginosa* and identify exploitable nutritional targets.

## Acknowledgments

The authors warmly thank Mamadou Godet and Sébastien Galopin for technical assistance, as well as Marie Leoz for submitting the transcriptomics results to the European Nucleotide Archive at EMBL-EBI.

## Author contributions

Caroline Martin-Duval, Conceptualization, Investigation, Writing – original draft, Writing – review and editing; Sandrine Dahyot, Conceptualization, Funding acquisition, Methodology, Supervision, Writing – review and editing; Erwan Bonnemains, Investigation; Enzo Debayle, Investigation; Abdellah Tebani and Soumeya Bekri, Metabolomic conceptualization, Formal analysis, Methodology, Review and Editing; Blandine Harel, Formal analysis, Methodology; Metabolomic conceptualization; Martine Pestel-Caron, Conceptualization, Funding acquisition, Methodology, Supervision, Validation, Writing – review and editing; Jean-Christophe Giard, Conceptualization, Investigation, Methodology, Supervision, Validation, Writing – original draft, Writing – review and editing.

## Data availability statement

The data that support the findings of this study, along with the supplementary tables, are openly available. Biofilm, survival, and metabolomics raw data and Supplementary Tables S1 and S2 (processed transcriptomic results derived from the RNA-seq analyses) are available in figshare at https://doi.org/10.6084/m9.figshare.32635260 under a CC BY 4.0 licence (46). Raw transcriptomics data are deposited in the European Nucleotide Archive (ENA) at EMBL-EBI under accession number PRJEB114131 (47).

This study is based on the research conducted as part of Caroline Martin-Duval’s PhD thesis entitled “*Μécanismes de survie et d’évοlutiοn adaptative de Ρseudοmοnas aeruginοsa dans l’arbre urinaire*”, which she defended in 2025 at the University of Rouen Normandy. The thesis is available at https://theses.hal.science/tel-05500505 (NNT: 2025NORMR089, tel-05500505) (48).

## Ethics approval

This study was conducted in accordance with the Declaration of Helsinki. Under the French Jardé Law (2016) and Article L. 1121-1 of the French Public Health Code, retrospective, non-interventional studies using anonymized, routinely collected data do not require specific informed consent or formal ethics approval with a permit number. The isolate collections were obtained as part of standard clinical care at the Rouen University Hospital, and all data were fully anonymized prior to analysis. The Ethics Committee for Research on Existing Data and/or Outside the Jardé Law at Rouen University Hospital, France, formally confirmed that no ethics approval was required for this study. Consequently, no reference or permit number was issued. The study was internally registered by the Directorate of Clinical Research and Innovation of Rouen University Hospital under the number 2018/413/OB.

## Supplemental material

Supplemental material is available online only.

